# Leaf traits predict water-use efficiency in U.S. Pacific Northwest grasslands under rain exclusion treatment

**DOI:** 10.1101/2022.09.23.509243

**Authors:** Hilary Rose Dawson, Toby M. Maxwell, Paul B. Reed, Scott D. Bridgham, Lucas C. R. Silva

## Abstract

Does drought stress in temperate grasslands alter the relationship between plant structure and function? Here we report data from an experiment focusing on growth form and species traits that affect the critical functions of water- and nutrient-use efficiency in prairie and pasture plant communities. A total of 139 individuals of 12 species (11 genera and four families) were sampled in replicated plots maintained for three years across a 520 km latitudinal gradient in the Pacific Northwest, USA. Rain exclusion did not alter the interspecific relationship between foliar traits and stoichiometry or intrinsic water-use efficiency. Rain exclusion reduced intrinsic water-use efficiency in grasses, an effect was primarily species-specific, although leaf morphology, life history strategy, and phylogenetic distance predicted intrinsic water-use efficiency for all twelve species when analyzed together. Variation in specific leaf area explained most of the variation in intrinsic water-use efficiency between different functional groups, with annual forbs and annual grasses at opposite ends of the resource-use spectrum. Our findings are consistent with expected trait-driven tradeoffs between productivity and resource-use efficiency, and provide insight into strategies for the sustainable use and conservation of temperate grasslands.

**Plain language summary:** Scientists have previously shown that plant leaf form (e.g., shape, width, size) has a predictable relationship to leaf function (e.g., how it can perform biological processes). When we deprive plants of water, does this relationship break down? We grew prairie and pasture plants at three sites in Oregon and Washington, USA, spanning a broad range of climate and water availability. At each site, we built shelters over half our plots to keep out some of the rain, reducing how much water our plants received. Leaf form-function relationships did not change between plots with more or less water. However, each species had a different water use efficiency and nutrient content, and some grasses had an unusual response, that is, they became less efficient at using water under less rain. Overall, we were pretty good at predicting water and nutrient use based on leaf form, whether plants were annual or perennial, and how related they were. Our findings match expectations about leaf structure-function relationships and people who manage temperate grasslands can use our results to decide which plants will work best for using and conserving their systems.

**Key points:**

- Foliar structure-function relationships did not change under experimental drought.
- Leaf morphology, life history, and phylogenetics predicted resource-use for 12 species.

## 1. Introduction

Grasslands in the western Pacific Northwest face an increased risk of drought stress due to rising temperatures, decreasing summer precipitation, and increasing evaporative demand (Dalton & Fleishman, 2021; Jung & Chang, 2012). Recent research suggests that drought stress has the potential to change grasslands, their species composition, and their forage production function (Mackie et al., 2019). Biodiverse communities may mitigate these negative changes, as functional differences between species, and related resource-use complementarity, provide a significant degree of resistance and resilience to drought stress (Hallett et al., 2014; Li et al., 2019). To better understand the mechanisms behind the stability documented in biodiverse grassland communities, and perhaps harness its benefits for conservation and management, we must consider the degree to which plants can morphologically and functionally adjust to persistent drought.

The relationship between plant functional traits and associated strategies for water and nutrient use provides a basis for interpreting the limits of stress response from individuals to communities and ecosystems (Enquist et al., 2015, 2017). For example, foliar morphology is related to primary productivity and nutrient cycling through a trait-dependent tradeoff termed the leaf economic spectrum (Wright et al., 2004). Notably, the same relationship that delineates the leaf economic spectrum, ranging from short-lived to long-lived leaves, with intrinsically high and low photosynthetic capacity respectively, makes morphological traits a proxy for potential foliar function such as the amount of carbon gained per unit of nutrient or water used for plant growth (Kröber et al., 2015). Indeed, the intrinsic water-use efficiency (iWUE; the ratio of net assimilation, A, to stomatal conductance, gs) and nutrient-use efficiency (e.g., reflected in leaf carbon to nitrogen ratios) of key functional groups tend to be correlated with plant growth form, life span, and leaf area per mass (i.e., specific leaf area, SLA). Although iWUE is not typically included in the leaf economic spectrum, it has been linked multiple times to traits that comprise the spectrum (Gouveia & Freitas, 2009; Soh et al., 2019; Temme et al., 2017). We can use these form-function relationships, measured at the level of individuals or species, to infer how stress responses might scale up to the level of communities and ecosystems, as in landscapes where variation in the relative cover of dominant species allows for leaf-to-canopy upscaling of water and nutrient costs of photosynthesis (Silva & Lambers, 2021). Click or tap here to enter text.

Here, we examined whether rain exclusion affects the foliar morphology, nitrogen and carbon to nitrogen ratios, and iWUE of 12 common grassland species at three sites spanning a 520 km latitudinal gradient in the Pacific Northwest. We examined how different functional groups responded to rain exclusion based on their growth strategies and trait plasticity. We focused of relationships that reflect differences in species life history strategy (annual or perennial) and functional group (grass or forb), both of which are expected to affect species reproduction and survival during drought stress (Tilman & El Haddi, 1992). Closely related species tend to resemble each other in form and function, a phenomenon known as “phylogenetic signal” (Blomberg & Garland, 2002). Because phylogenetic signal is not random across a subset of species, it is important to include phylogeny as a variable when testing patterns of drought stress acclimation to control for the “relatedness” of species of interest (Avise & Wollenberg, 2010). To study form-function relationships we also focused on phylogenetic differences between species influenced how traits varied in response to drought. Specifically, we used a climate gradient, from cool and wet to warm and dry, to test the hypothesis that differences in species’ sensitivity to drought would be explained by their physiology (leaf traits, life history strategies, and functional group) and the phylogenetic distance between them. Our experiments provide insight to the physiological and environmental mechanisms that link species form and function, which could help improve temperate grassland management and conservation.

## 2. Materials and Methods

### 2.1 Site descriptions

We conducted replicated experiments at three different sites along a 520 km latitudinal gradient in the Pacific Northwest (Fig. 1, Table 1). The study areas share a Mediterranean climate with increasingly hotter and drier summers from north to south, where we see an earlier onset of summer senescence, despite the southern site having the highest mean annual precipitation (Table 1). Each experimental site contained 20 plots: 10 had their species composition manipulated as part of separate phenology and demography experiments (Reed, Bridgham, et al., 2021; Reed et al., 2019), and 10 had their species composition unmanipulated and consisted primarily of the already established pasture grasses that dominated at each site prior to the experiment. The manipulated plots were mowed, raked, received herbicide, and seeded with a mix of 29 native prairie grass and forb species between 2014-2015, followed by repeated seeding with 14 native grasses and forbs in fall 2015, 2016, and 2017 (Reed et al., 2019), a process that is analogous to typical restoration efforts in the region.

**Figure 1.**
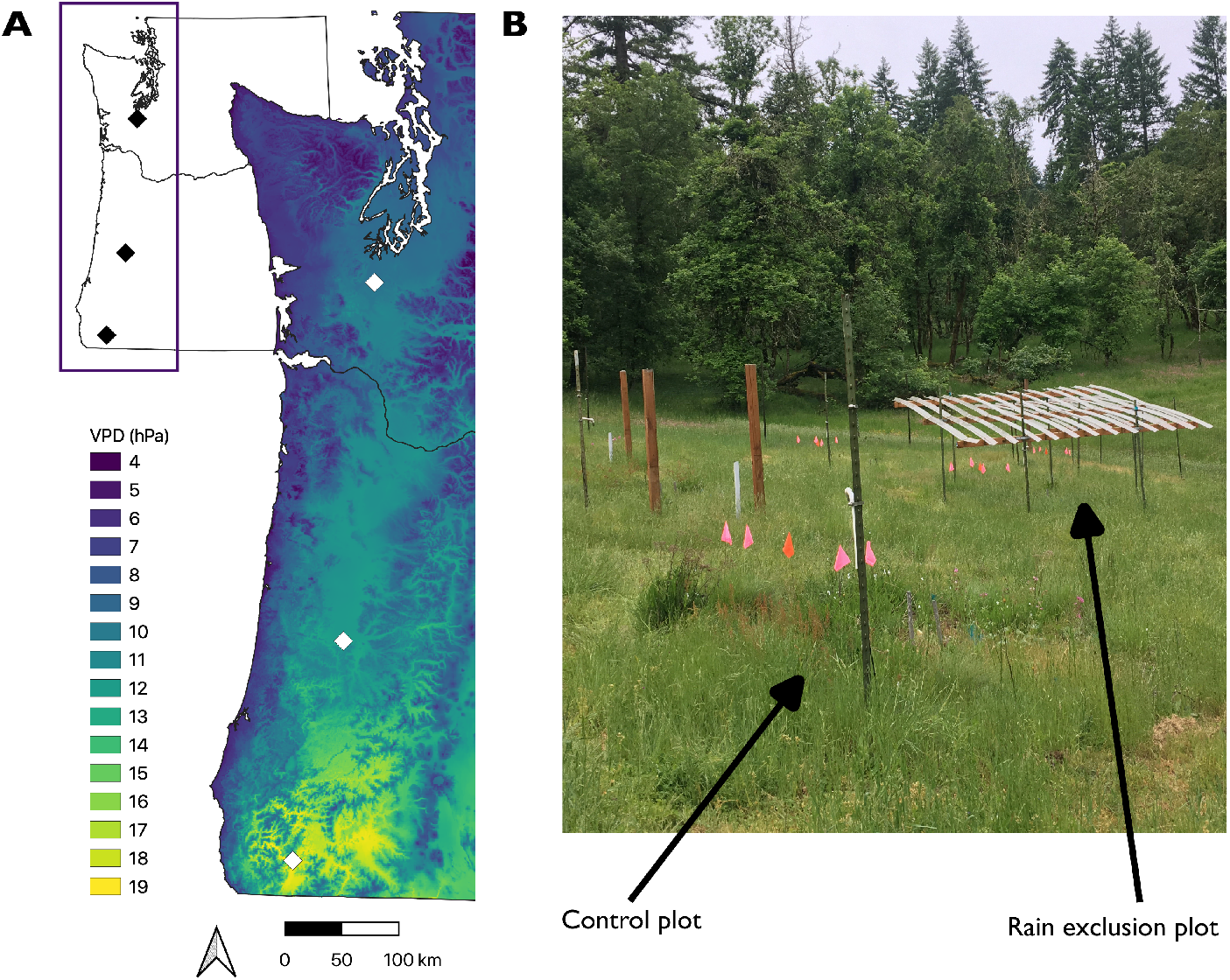
a. Site locations with maximum vapor-pressure deficit (VPD) expressed as hectopascal (hPa), or 100 x 1 pascal, pressure units which are equal to millibar pressure unit, and b. example of experimental set up depicting a rain exclusion shelter. VPD data is from PRISM for 1991 to 2020 (http://www.prism.oregonstate.edu/normals/). Points show site locations in Washington and Oregon.

**Table 1.** Characteristics of the three experimental site locations.

|  | Experimental Site |  |  |
| --- | --- | --- | --- |
|  | Southern | Central | Northern |
|  | Selma, Oregon | Eugene, Oregon | Tenino, Washington |
| Latitude; longitude | 42.27811; -123.642278 | 44.02615; -123.182171 | 46.86415; -122.958918 |
| Elevation (m) | 394 | 165 | 79 |
| Mean air T (°C) | 10.26 ± 6.80 | 10.70 ± 6.49 | 8.68 ± 5.52 |
| Min air T (°C) | 2.11 ± 4.56 | 5.49 ± 4.89 | 2.43 ± 4.69 |
| Max air T (°C) | 20.10 ± 9.76 | 17.63 ± 8.88 | 15.25 ± 7.61 |
| Mean annual precipitation (mm) | 1325 ± 327 | 894 ± 207 | 1300 ± 247 |
| Soil taxonomy | Loamy-skeletal, mixed, superactive, mesic Entic Ultic Haploxerolls | Very-fine, smectitic, mesic Vertic Haploxerolls | Medial, mixed, mesic Typic Haploxerands |

Notes: Climate data calculated from site specific dataloggers in 2017 and 2018. Minimum and maximum calculated as annual means of daily averages. Errors represent standard deviation. Elevation and soil taxonomy from Reed et al. (2019).

For both the manipulated and unmanipulated plots, five were randomly assigned to a rain exclusion treatment while the remaining five were assigned as controls. We estimate to have experimentally imposed reduced annual rainfall by ~40% using a fixed rainout shelter design blocking no more than ~8% light transmittance (Yahdjian & Sala, 2002). Rainout shelters were erected in February 2016 for the manipulated plots and February-March 2017 for the unmanipulated plots and maintained until after our sampling in summer 2019 (see Fig. S1 and S2 for the effects of the rainout shelters; see Supplemental Data with this article). The vegetation plots under the shelters were circular with 1 m diameter. Each plot was nested within a larger 3 m diameter plot. Rainout shelters were 3.7 m x 3.7 m squares and stood 1.5 m above the vegetation plots, sloped to 1 m above the plot on the far side to promote drainage, providing ~30 cm buffer around the vegetation plots. Phenology and demography information from these experimental plots have been previously published in (Peterson et al., 2021; Reed, Bridgham, et al., 2021; Reed et al., 2019; Reed, Pfeifer-Meister, et al., 2021).

### 2.2 Species and phylogenetic data

For this study, we selected 12 species that were abundant in at least one site (Fig. 2, Table S1). Click or tap here to enter text.The species studied represented a total of 139 individuals spanning 11 genera and four families. Two species (*Bromus hordeaceus* and *Sidalcea malviflora* ssp*. virgata*) were present across all three sites; the former was present in both manipulated and unmanipulated plots across all sites. We collected up to three leaves for each species in each plot. Because not all species were present at all sites or all plots, our sampling was uneven; however, this unevenness was distributed throughout the relevant groups and should have biased our analyses (Table S2). Leaves were sampled on 8 May 2019 (southern site), 29 May 2019 (central site), and 6 June 2019 (northern site) at approximately peak growing season in each site. For each species at each plot, leaves were taken from multiple mature individuals where available and were stored in envelopes in paper bags which were dried for at least 48 hours at 50°C before being stored at ambient conditions. All species (including all grasses) included in this study are described as possessing cool-season (C3) metabolism (Jackson et al., 2010; Osborne et al., 2014).

**Figure 2.**
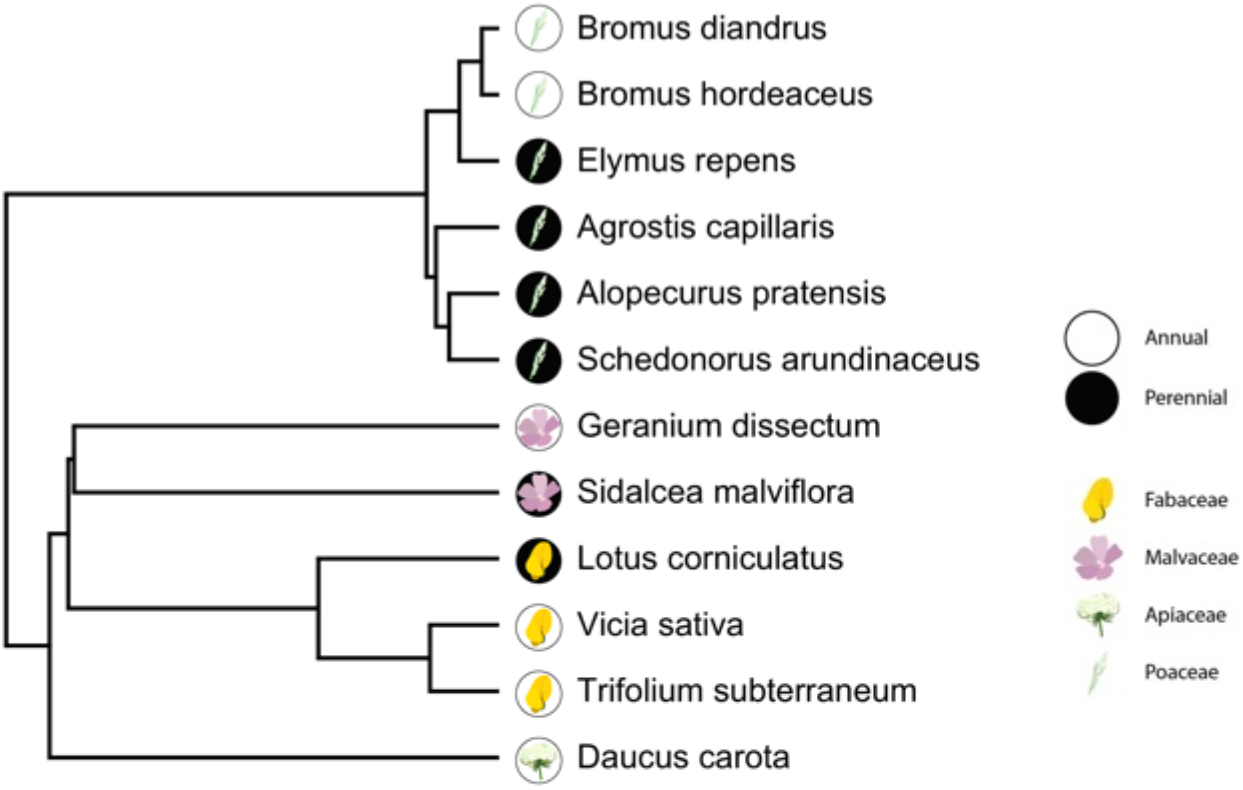
Phylogeny of the twelve species from four families that were examined in our data. Phylogenetic data and tree structure is pruned from the mega-tree in Jin and Qian (2019). Symbols represent family membership with backgrounds that show life history.

We determined plant functional group based on family where members of Poaceae were designated ‘grasses’ and all other plants (all non-graminoid) were designated ‘forbs’. We used the USDA PLANTS database to establish life history strategy (USDA NRCS 2019). This database designated three species (*Bromus diandrus, Daucus carota*, and *Geranium dissectum*) as either annual/biennial or biennial. We treated these species as annuals, reflecting their shorter lifespan compared to ‘true perennials’.

All analyses were conducted in R v.3.6.1 (R Development Core Team, 2017). To make it easy to reproduce our analyses, we use the format of R package::function, followed by the appropriate citation for each package mentioned. Phylogenetic data, including tree structure and branch lengths, were derived from a mega-tree compiled by Jin and Qian (2019) using two well-established, recent mega-trees based on molecular data and the Angiosperm Phylogeny Group 2016 (Chase et al., 2016). The tree was pruned with V.PhyloMaker::phylo.maker and the resulting tree was both ultrametric and binary. To integrate phylogenetic distance into our traits dataset, we calculated mean phylogenetic distance (also known as mean pairwise distance) using a method modified from Ness et al. (2011). Using this method, we created a cophenetic table using stats::cophenetic. From this table we calculated mean phylogenetic distance per species, a metric which we defined as the mean of the phylogenetic distance between a set species and every other species in a cophenetic table.

### 2.3 Leaf trait data

To measure leaf area, we scanned one leaf sample per individual at a standardized cropping area at 600 dpi (see table S1 for leaves per species). We prepared images with the magick package (Ooms, 2019) and analyzed leaf area with LeafArea::run.ij (Katabuchi & Masatoshi, 2015; Rueden et al., 2017). We weighed each sample on a Sartorius CP2P-F microbalance (Göttingen, Germany) and calculated specific leaf area (SLA) as area divided by mass. We used SLA from dried material to test relationships with nutrient content and iWUE, inferred from stable carbon isotope ratios, assuming that SLA from dried material scales to SLA measured with fresh material (Perez & Heberling, 2020).

Stable isotope analysis was conducted by the Stable Isotope Facility at the University of California, Davis, on a PDZ Europa ANCA-GSL elemental analyzer interfaced to a PDZ Europa 20-20 isotope ratio mass spectrometer (Sercon Ltd., Cheshire, UK). We calculated iWUE according to Werner et al. (2012). The isotopic discrimination compared to the atmosphere (Δ^13^*C*) was calculated in Eq. 1 using the δ^13^C values of the air (*δ*^13^*C_a_*) derived from Keeling et al. (2010) and the measured plant (*δ*^13^*C_p_*):

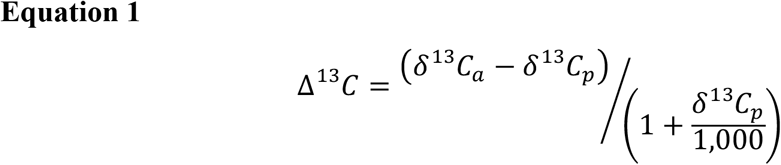

We used the isotopic discrimination in Eq. 1 and the diffusive and photosynthetic fractionation constants (a: 4.4‰ and b: 28‰) to calculate the ratio of CO_2_ partial pressures inside the leaf (C_i_) and in the atmosphere (C_a_).

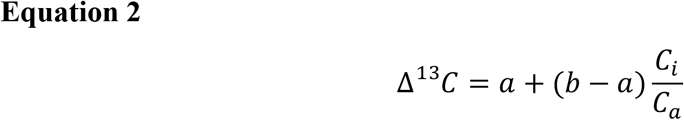

From this, we solved for iWUE, scaling to 1.53 (the ratio of water vapor to CO2 diffusivity).

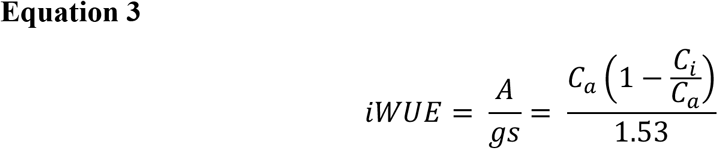

Leaf nitrogen (N) was collected as part of the stable isotope analysis. Leaf carbon to nitrogen ratio (C:N) was calculated as

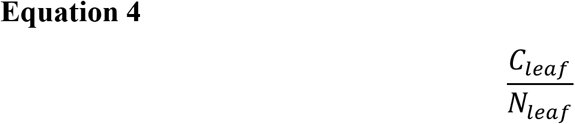

### 2.4 Site data

Since the three experimental sites have unique soil characteristics, we calculated soil matric potentials as a comparable metric of plant moisture availability. Daily matric potentials were calculated from daily volumetric water contents and site-level measurements of percent sand, clay, and soil carbon using methods described in Saxton & Rawls (2006). We aggregated data to daily values before calculating growing season values, and we defined a growing season as October 1 to June 30, beginning with the return of fall rains and ending with the onset of summer drought (fig 3; Reed et al., 2019). We collected six soil cores from each site, dried the soil for 48 hours at 60°C and sieved it to 2 mm. We then determined percent sand from the weight after sieving to 53 μm, percent clay using the hydrometer method (Gee & Bauder, 1986), and total soil carbon using a Costech Analytical Technologies 4010 elemental combustion analyzer (Valencia, CA, USA).To correct the left-skewed data, we log-transformed for a more normal distribution. We used -log(1-SMP) units to be consistent with raw values where negative values indicate drier conditions. Soil temperature were measured in situ and continuously logged at all three sites along with daily volumetric water content.

**Figure 3.**
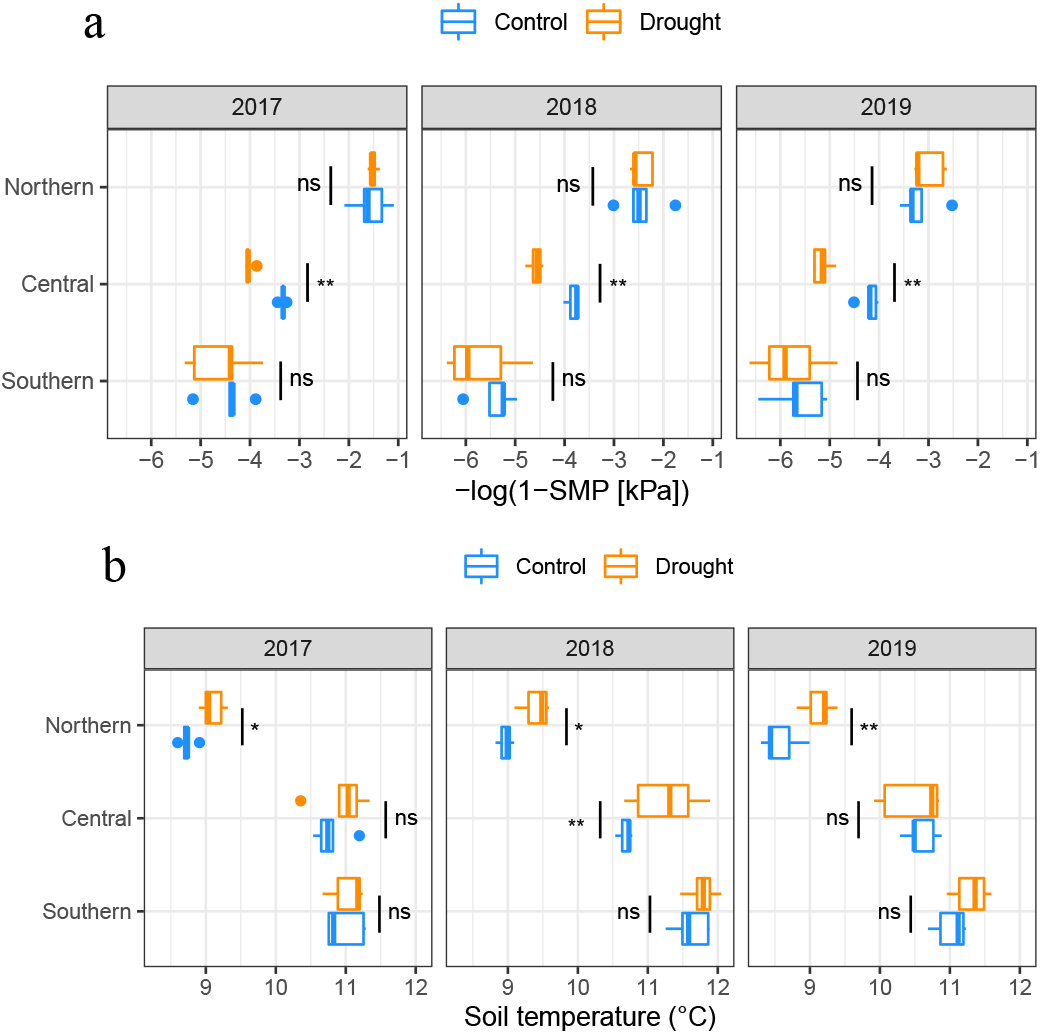
Effects of rain exclusion treatment on a) soil matric potential and b) soil temperature at each site for each growing season. Growing season is defined as 1 Oct to 30 Jun and each season is labeled by the spring months of the growing year (e.g. 2018 is 1 Oct 2017 to 30 Jun 2018). Asterisks indicate significant differences tested by ANOVA (ns = not significant, ** = 0.001, *** < 0.001, see table S5 for results and table S7 for summary statistics and figs S1 and S2 for daily variation). Median values are indicated by the central vertical line, with vertical lines to the left and right indicating the interquartile range. Horizontal lines indicate the interquartile minimum and maximum. Dots indicate outliers.

### 2.5 Statistical analyses

We tested the relationships between functional and structural traits using linear correlations and mixed-effects models. The measured traits (SLA, iWUE, C:N, and N) were not normally distributed, so we log-transformed the data prior to performing statistical analysis. We determined plant growth strategy, site, and treatment effects on each trait using Type II analysis of variance (ANOVA), including interaction terms. We calculated post-hoc significance using Tukey-adjusted P-values. Response ratios were calculated as

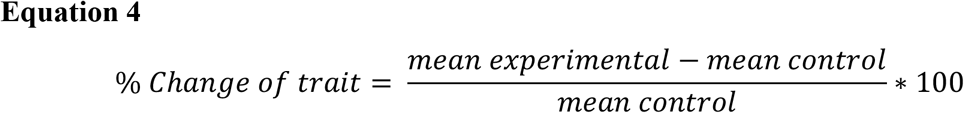

by species in each manipulation treatment at each site (Myers et al., 2014).

We ran ANOVAs for each growing year to determine if the rain exclusion treatment had a significant effect on soil matric potential. The interaction between site and drought treatment (term site x treatment) was significant, so we ran t-tests within each site to see where rain exclusion was significant.

At each site, we calculated means and standard errors of each trait for each species and used these data to determine phylogenetic signal (i.e. the strength of phylogeny on species’ traits) using phytools::phylosig, using Pagel’s λ. We used mean pairwise distance to test the phylogenetic signal across our entire dataset, and we used that signal as one of the predictive variables in a mixed effect model, effectively including it as a constraint on species responses to drought (De Vienne et al., 2011).

We then constructed a principal component analysis (PCA; Fig. S3) of the four continuous trait datasets (iWUE, SLA, C:N, and N) using stats::prcomp. We hypothesized seven biologically sound mixed effect models and determined the strongest with Akaike’s information criteria with small sample bias adjustment (AICc) using AICcmodavg::aictab (table S3; Mazerolle, 2006, 2020). Some degree of multicollinearity was expected in the mixed-effects models due to associations between structural and functional leaf traits, which are inherently correlated. However, the PCA analysis shows a near orthogonal (i.e. perpendicular) trait continuums: iWUE-SLA and C:N-N which suggests independent contribution of those variables to mixed-effects models. All models excluded the functional group effects shown in the response ratio analysis because functional groups represent categorical variables that summarizes phylogenetic differences included in the mixed models as a continuous variable (i.e., phylogenetic distances). Errors are shown as standard deviation unless specified otherwise.

## 3. Results

### 3.1 Overall effects of rain exclusion

Rainout shelters reduced soil matric potential in the central site in all years but had a limited effect in the other two sites (Table S4, Fig. S1). Rainout shelters also increased soil temperatures in the northern site in all three years and in the central site in 2018 (Table S5, Fig. S2).

Response ratios of rain exclusion compared to control plots showed that rain exclusion had a significant effect on only annual and perennial grass iWUE at 95% confidence interval at all sites (Table 2). Specifically, the rain exclusion treatment significantly reduced grass iWUE compared to the control but had no significant effects on the other measured functional forbs or group or any other grass traits (Table 2, Fig. 4). Site and the interaction term between site and rain exclusion treatment were not significant for all variables except SLA (Table 2), which was highest in the north (0.154 ±0 .05 cm^2^/mg) and lowest in the south (0.130 ± 0.04 cm^2^/mg).

**Table 2.** Statistical summary of four full factorial mixed effect Type II ANOVAs testing the effects of rain exclusion treatment and sites on functional species traits.

| Predictor |  | log(iWUE) |  | log(SLA) |  | log(C:N) |  | log(N) |  |
| --- | --- | --- | --- | --- | --- | --- | --- | --- | --- |
| *Fixed effects* | *df* | $\chi^2$ | *P* | $\chi^2$ | *P* | $\chi^2$ | *P* | $\chi^2$ | *P* |
| **Treatment** | 1 | 4.019 | **0.045** | 0.266 | 0.61 | 0.590 | 0.442 | 0.840 | 0.359 |
| **Functional group** | 1 | 55.834 | **<0.001** | 38.660 | **<0.001** | 17.698 | **<0.001** | 13.466 | **<0.001** |
| **Life history strategy** | 1 | 41.405 | **<0.001** | 179.915 | **<0.001** | 6.794 | **0.009** | 4.582 | **0.032** |
| **Site** | 2 | 1.605 | 0.448 | 35.625 | **<0.001** | 2.277 | 0.320 | 1.183 | 0.553 |
| **Treatment:Functional group** | 1 | 0.295 | 0.587 | 0.717 | 0.40 | 0.341 | 0.559 | 0.393 | 0.531 |
| **Treatment:Life history strategy** | 1 | 0.374 | 0.541 | 2.071 | 0.15 | 2.851 | 0.091 | 2.468 | 0.116 |
| **Life history strategy:Functional group** | 1 | 7.940 | **0.005** | 12.885 | **<0.001** | 2.967 | 0.085 | 2.144 | 0.143 |
| **Treatment:Life history strategy:Functional group** | 2 | 0.852 | 0.653 | 0.754 | 0.69 | 1.766 | 0.414 | 2.083 | 0.353 |
| **Treatment:Site** | 1 | 0.002 | 0.968 | 0.206 | 0.65 | 0.547 | 0.460 | 0.687 | 0.407 |
| *Random effect* |  |  |  |  |  |  |  |  |  |
| **Plot** | 1 |  | 1 |  | 0.839 |  | 0.248 |  | 0.236 |

**Figure 4.**
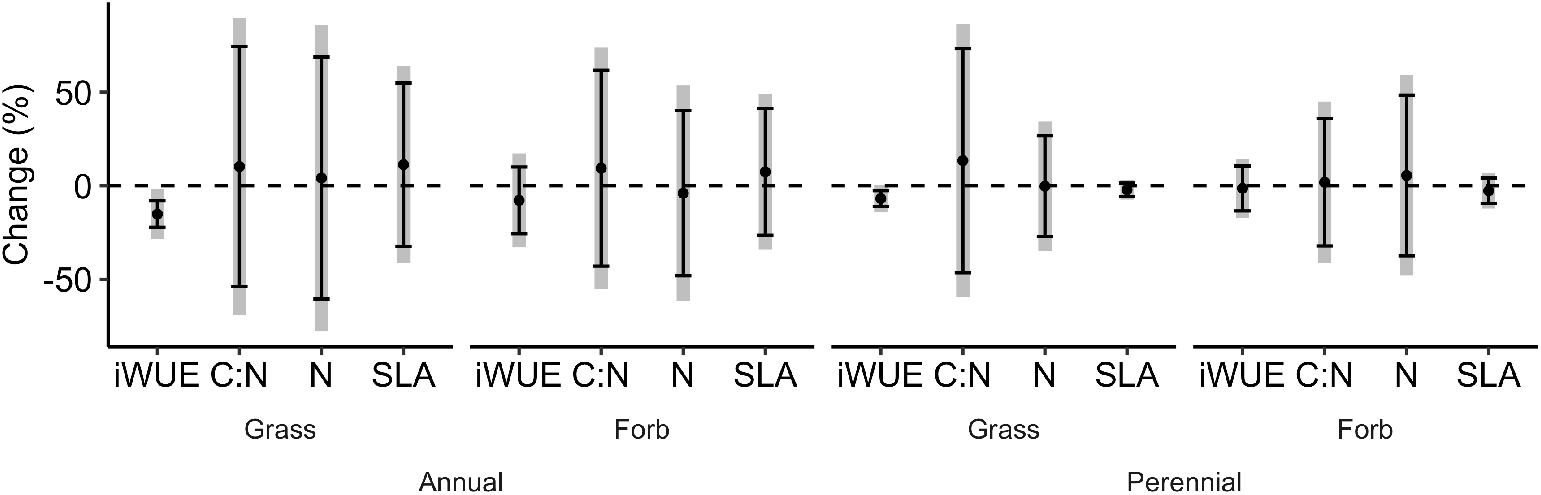
Response ratios measured as percentage change in traits at 40% reduced rainfall relative to unaltered rainfall. There was a significant interaction between life history strategy and functional group for iWUE and SLA (Table 2). Error bars represent 95% confidence intervals. Grey bars represent 90% confidence intervals. Traits include intrinsic water-use efficiency (iWUE), carbon to nitrogen ratio (C:N), nitrogen content (N), and specific leaf area (SLA).

### 3.2 Predicting intrinsic water-use efficiency

The strongest statistical model to predict iWUE from all measured species traits and environmental variables included SLA, mean phylogenetic distance, C:N, N, life history strategy, and rain exclusion treatment (Fig. 5, Table S3). Variation in SLA explained most of the variation in iWUE between different functional groups, with annual forbs and annual grasses at opposite ends of the resource-use spectrum. Notably, functional group and life history strategy significantly affected iWUE, SLA, C:N, and N content (Table 2). However, each variable divided into two nearly orthogonal trait continuums: iWUE-SLA and C:N-N, continuums which had a nearly orthogonal relationship that explained 86.6% of variation across species and sites (Fig. S3). Life history strategy was divided along the iWUE-SLA continuum, where annuals were associated with higher SLA and perennials with higher iWUE. Nutrient status assessed as leaf C:N and total N content had the least effect of the factors in the strongest mixed-effects model.

**Figure 5.**
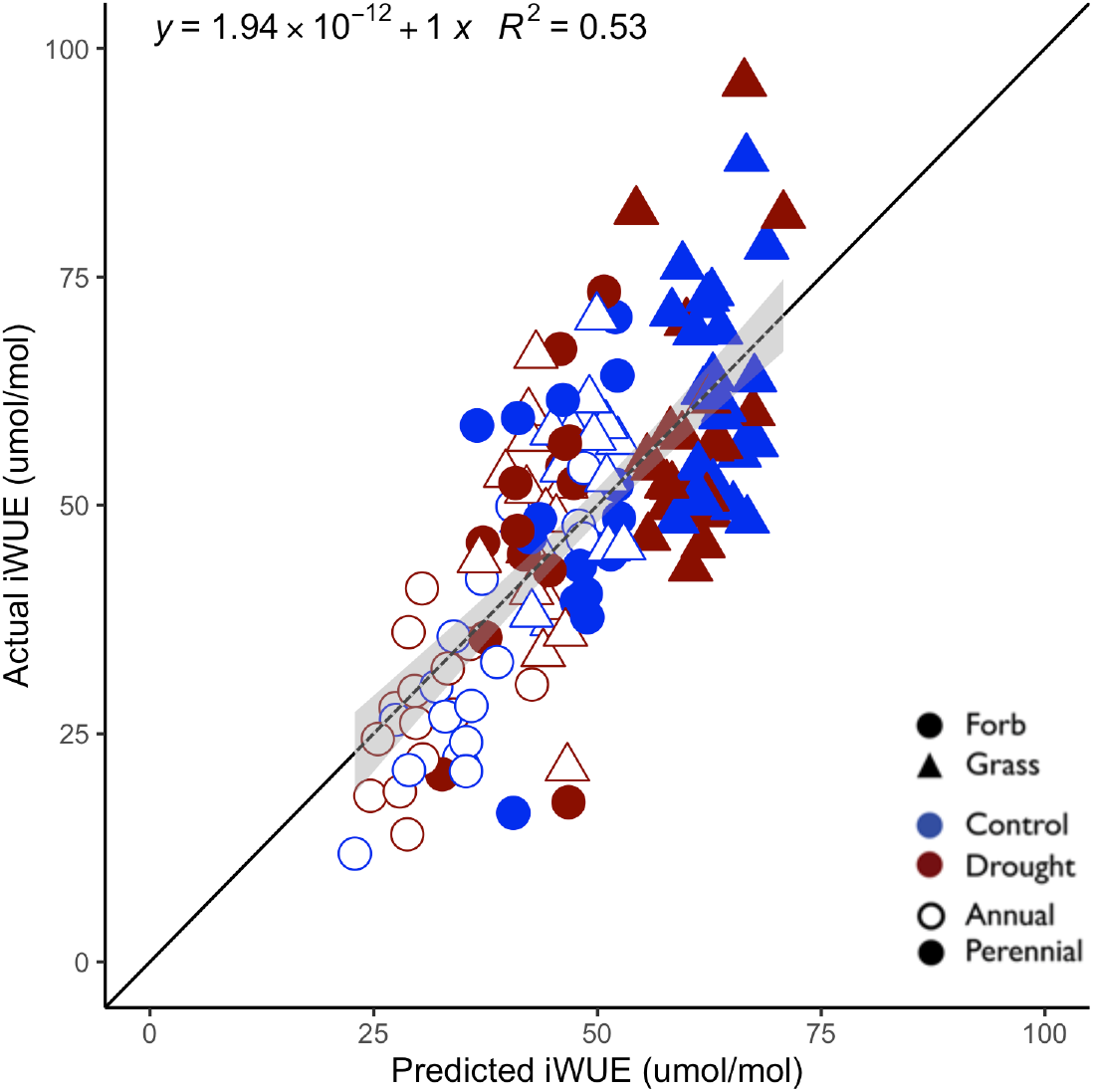
Statistical model of intrinsic water-use efficiency (iWUE) observed versus predicted by species traits and environmental variables. Black line indicates 1:1 relationship. Grey line indicates linear relationship described by equation in top left. Light grey indicates standard error. There was a significant interaction between life history strategy and functional group (Table 2). The significance of the model is detailed in Table S8.

### 3.3 Model variables and interactions

There was a strong inverse relationship between iWUE and SLA (Fig. 6, R^2^ = 0.67, P < 0.001), but SLA had no significant effect with C:N or N (Fig. S5). The structure-function relationship follow expectations for differences between major functional groups, as reflected in significant effects of life history strategy (annual or perennial) and functional group (grass or forb) on iWUE and SLA but not for C:N or N. Beyond functional categories, iWUE had a significant association with continuous phylogenetic distances between species (Pagel’s λ = 0.78, P = 0.038) but SLA (λ = 0.18, P = 1), C:N (λ = 0.14, P = 0.78), and N (λ < 0.30, P = 1) did not.

**Figure 6.**
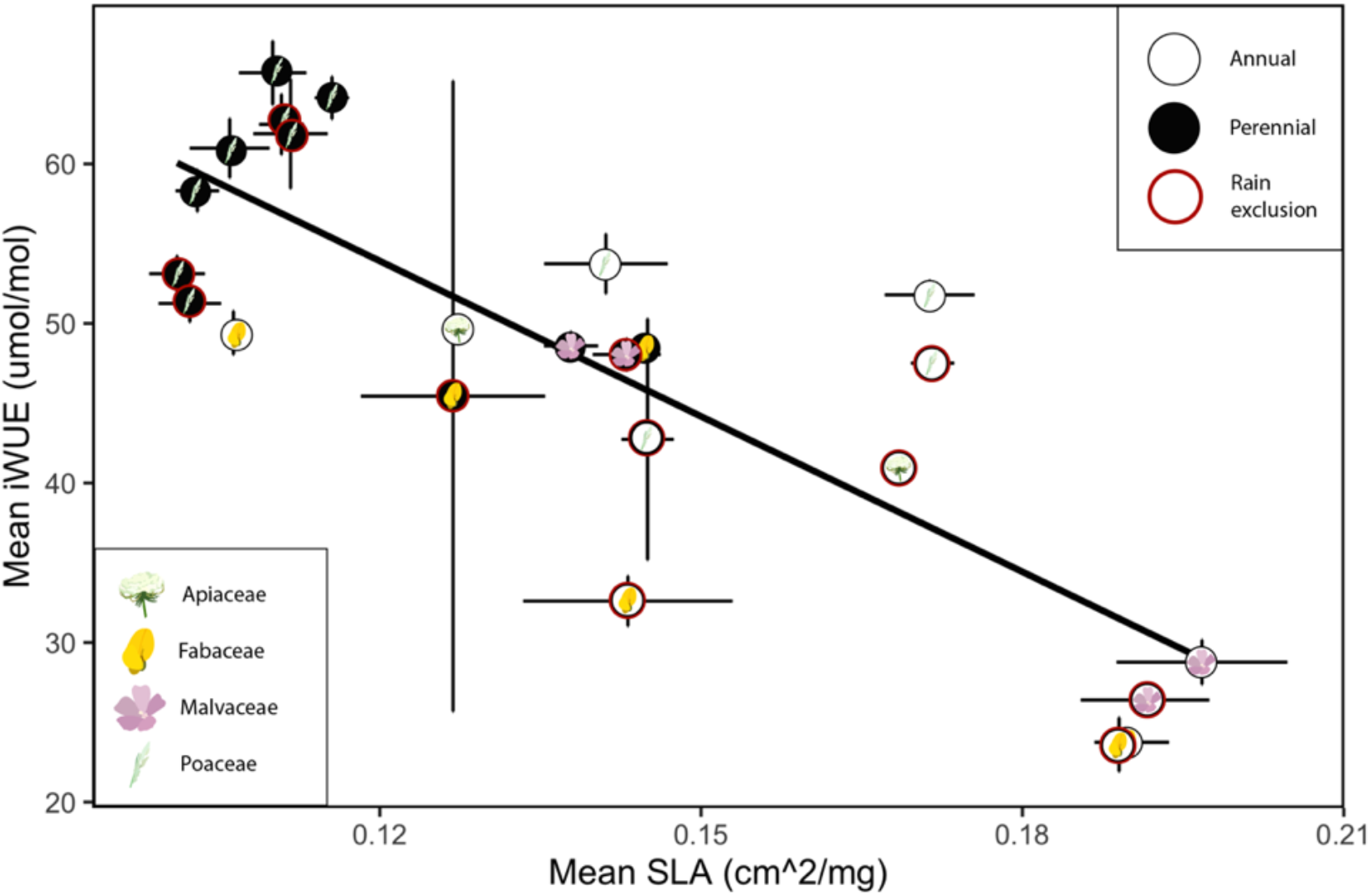
Linear correlation plot of mean intrinsic water-use efficiency (iWUE) with mean specific leaf area (SLA). Each point represents the mean value by species and rain exclusion treatment. There was a significant interaction between life history strategy and functional group; Table 2). Plot of all data points is available in Fig. S4. Symbols represent different families. Symbol fill represents life history strategy (black fill = perennial, white fill = annual). Red outline indicates the rain exclusion treatment. Fitted line is significant (R^2^ = 0.67, P <0.0001, y = −324.71x + 92.79). Errors indicate standard deviation.

Despite a significant interaction between life history strategy and functional group, on average, perennials had lower SLA and higher iWUE compared to annuals (Tables 2 and 3). In particular, perennial grasses had the lowest observed SLA (0.11cm/mg^2^ ± 0.24 SD) while annuals (both grasses and forbs) had the highest SLA (0.16 cm/mg^2^ ± 0.04 SD and 0.18 cm/mg^2^ ± 0.05 SD respectively; Tables 2 and 3). In most cases, differences in SLA translated into differences in iWUE, with perennial grasses showing the highest iWUE (60.38 μmol/mol ± 11.93 SD) while annual forbs had the lowest iWUE (29.82 μmol/mol ± 10.41 SD). Life history strategies and growth forms were not significantly divided across the continuum of plant N content. However, perennial forbs had significantly lower C:N than other functional groups and forbs had higher leaf N concentration than grasses but not significantly so.

**Table 3.** Summary of mean trait values by life history strategy and functional group.

|  | Annual |  | Perennial |  |
| --- | --- | --- | --- | --- |
| Trait | Grass | Forb | Grass | Forb |
| SLA (cm/mg<sup>2</sup>) | 0.16 ± 0.03 *a* | 0.18 ± 0.04 *a* | 0.11 ± 0.02 *c* | 0.14 ± 0.04 *b* |
| iWUE (μmol/mol) | 49.57 ± 10.60 *b* | 29.82 ± 10.41 *c* | 60.38 ± 11.93 *a* | 48.17 ± 13.94 *b* |
| C:N | 24.05 ± 6.47 *a* | 22.73 ± 11.82 *a* | 23.90 ± 10.91 *a* | 16.33 ± 4.87 *b* |
| N (%) | 1.87 ± 0.49 | 2.30 ± 1.05 | 2.01 ± 0.70 | 2.70 ± 0.72 |

Notes: Letters indicate post-hoc (Tukey's honestly significant difference) significant differences for each trait among the four life history strategy and functional group combinations that were statistically different (see Table 2). Traits include intrinsic water-use efficiency (iWUE), carbon to nitrogen ratio (C:N), nitrogen content (N), and specific leaf area (SLA). Errors indicate standard deviation.

## 4. Discussion

Our data show that leaf traits and stoichiometry predicted iWUE across a range of species under rain exclusion and control plots along a broad latitudinal Mediterranean-climate gradient. Our rain exclusion treatment impacted iWUE in grasses (the only trait and functional group affected by treatment; Table 2), although in an unexpected direction. Previous studies using similar rain shelters have found iWUE increases under drought conditions (e.g., Beer et al., 2009; Ocheltree et al., 2020); however, in our study the grasses (particularly the annual grasses) had significantly reduced iWUE in the rain exclusion treatment compared to controls (Fig. 4). This could be an effect of phenology where flowering plants have greater iWUE than plants in which flowering is delayed due to reduced rainfall (Franks, 2011); however, a different study previously conducted at the same experimental sites did not show any significant changes in phenology in the rain exclusion treatment (Reed et al., 2019).

Most annual grasses in this system (and indeed, those in our study) are nonnative, winter-growing species that are increasing in abundance with warming (Reed, Pfeifer-Meister, et al., 2021). These species seem to have an avoidance mechanism to drought by their winter-growing, early-maturing strategy and are able to outcompete other functional groups, perhaps explaining a reduced iWUE, as inferred from the isotopic signal which captures the integrated effect of environmental conditions (e.g. VPD, soil water, and air temperature) representative of the whole leaf lifespan (Maxwell et al., 2018). Change in phenology could therefore lead to unexpected shifts in iWUE by reflecting the less stressful period of leaf development for species that grow fast and early in the season to avoid drought stress, such as nonnative grasses. Additionally, the relative cover of such drought-adapted species likely affected the performance of the individuals measured here without changing the general trend between form and function across species (Figs. 4 and 5). For example, the rain exclusion treatment only had an effect in the shoulder fall and spring seasons because the Mediterranean climate contrasts very wet winters with dry summers, as shown in soil matric potential across the sites and treatments (Fig. S1). Further, the seasonality and absolute amount of available soil water, as represented by a natural climate gradient and a rain exclusion treatment, had relatively minor effects on iWUE in a Mediterranean-type climate.

Soil type and site-specific microclimate are also important controls of plant physiological performance, and thus water and nutrient use, and yet we see a consistent and predictable relationship between form and function across species at all sites and experimental treatments. Indeed, despite differences in soil type and microclimate, we found a strong relationship between iWUE and SLA (Fig. 6). Although the leaf economic spectrum does not typically include iWUE, this structure-function relationship holds across species with a range of nitrogen-use strategies, but is not consistent within related taxa, such as Poaceae.

### 4.1 Stoichiometric relationships

Plant growth strategies (annual vs. perennial, grasses vs. forbs) had statistically significant effects on leaf functional traits (Table 3). On average, perennials were associated with increased iWUE and decreased SLA compared to annuals. However, there was a significant interaction term between functional group and life history group, whereas life history strategies and growth forms did not significantly correlate with plant N content. The relationships between iWUE-SLA and C:N-N were expected to be co-linear, but they formed two quasi orthogonal axes (Fig. S3). Evidently, C:N and N were strong covariates (Fig. S4), but we included both in the statistical model because there is evidence that each may respond differently to environmental stressors (Li et al., 2015), and thus represent potentially distinct ecological processes. In particular, our results concur with Li et al.’s (2015) findings that iWUE is not correlated with N or C:N. Excluding either N or C:N marginally raised the power of the PCA to 87% (data not shown) but this did not adequately show the distribution of the data because C:N had a weaker relationship with SLA compared to iWUE (Fig. S4). We suspect that this is due to differences in symbiotic and competitive interactions that occur belowground and that shape the differential investments in leaf structure and resource use within and across functional groups, as discussed above for nonnative grasses. Although all of our focal species participate in arbuscular mycorrhizal interactions (Table S1), only three species (all members of Fabaceae) have the capacity to fix N. The large variation around the mean within and across species could represent differences in phenology as well as variation in root development and common mycorrhizal networks, which were not measured in this study. This variation would eventually be reflected in the integrated measure of ecophysiological performance and should be a research priority going forward. In all but one instance, nitrogen-fixing species displayed lower than average iWUE relative to the general trend expected from variation in leaf area (Fig 5). This could be explained by the greater evaporative loss observed in other Fabaceae species (e.g. alfalfa) which make them outliers in terms of evaporative enrichment. For example, we have previously observed this phenomenon in oxygen isotope signatures of cellulose and bulk lipid extracts in field experiments where the isotopic signatures of the source soil water could be compared with the leaf water evaporative enrichment across multiple collocated functional groups (Silva et al., 2015).

### 4.2 Rain exclusion and water availability

Our sites span ~520 kilometers in latitude with a strong drought gradient where drought has an earlier spring onset at the southern sites; however, site did not affect iWUE (Table 2). Our data show that changing the amount of precipitation in this Mediterranean climate gradient had a small effect on iWUE, an effect that is unexpected due to the known link between iWUE and drought stress (e.g., Farquhar et al. 1989) and deserves further investigation in future studies. The effects of the rain exclusion treatment were species-specific and less detectable when analyzing species by common groupings (Fig. 4), although leaf traits varied according to life history strategy and functional group (Table 3). Clark et al. (2012) found that plant traits predicted bunchgrass prairie restoration success in the Pacific Northwest equally well as individual species status. However, they analyzed a suite of traits with greater breadth and depth, and they found that the effect varied significantly between sites. While measuring iWUE typically requires access to expensive equipment, we found that it can be reliably inferred from SLA, mean phylogenetic distance, and life history strategy (Fig. 5, Table S8). These are all metrics that can be measured with simple equipment (SLA) or determined using widely available information (mean phylogenetic distance, life history strategy).

### 4.3 Phylogenetic relationships

Overall, the generalizable leaf trait relationships that we found in our data allowed us to identify tradeoffs in plant resource-use for different growth forms. These relationships can act as a new lens for land managers who can use these relationships to target species when managing Pacific Northwest grasslands. This could manifest as a tool to aid in designing seed mixes for functional restoration in a particular site or region. New perspectives such as this are critical for adaptive management that may also mitigate the impacts of climate change. Structure-function relationships can be used to improve on the relatively coarse restoration paradigm that biodiversity alone is the key to resistance and resilience to future climates. For example, we found that species relatedness measured as phylogenetic distance improved the strength of our model when used with functional traits (Table S3). Phylogenetic data are now widely available and accessible with open source software such as the V.PhyloMaker R package (Jin & Qian, 2019). Although there are multiple functions that can incorporate and analyze single trait values for each leaf of a phylogenetic tree, mean phylogenetic distance is different in that it can be used as a standalone trait in statistical models. It has been successfully adapted to measure the relatedness of native and exotic organisms, testing correlation between an organism’s relatedness to native species and predicting species introduction success (Ness et al., 2011; Van Wilgen & Richardson, 2011). However, it is worth noting that key morphological traits such as SLA was predicted by the categorical grass/forb designation but not the continuous phylogenetic distance variable. Although our study only includes twelve species, our results suggest that phylogenetic analysis when coupled with trait-based analysis could become valuable in ecological predictions.

### 4.4 Experimental limitations

There are caveats to our findings. We only installed plot-specific temperature and soil moisture dataloggers in manipulated plots so our localized climate data were limited (Table S1). Additionally, as explained above, the rain exclusion treatment only had a significant effect on soil water content across the entire growing season at the central site. During the dry summer months, there is negligible precipitation so a 40% reduction is not biologically meaningful. As a result, at least at the southern and northern sites, there may only be short shoulder seasons during the early fall and late spring when a reduction in rainfall has a measurable effect on soil water content. To the extent that the shelters had an effect, they would reduce evapotranspiration, which would warm the soil (Fig. S2). This suggests that the shelters were more effective than shown by the soil moisture data. Also, previous studies at the same site have found that soil moisture and plant community composition affect each other, potentially obscuring the effects of the rainout shelters(Reed, Pfeifer-Meister, et al., 2021). The results show that iWUE varies primarily with functional group (grass vs. forb) and life history (annual vs. perennial), with marked differences between nitrogen-fixer (i.e. Fabaceae) consistent falling below the average iWUE as predicted by leaf structure, as explained above (Fig 5).

The weak effects of the rain shelters, combined with the relatively low number of sampled individuals for each of the 12 species, makes it difficult to assert whether variation in precipitation would alter intraspecific traits as expected. Additionally, we calculated iWUE from δ^13^C values, a proxy that is well-established but can be blurred by compounding factors unrelated to actual water use. One of the most notable of these is variation in phenology (i.e. leaf age) on mesophyll conductance which can act on ^13^C conductance without impacting iWUE. The equation for iWUE can be scaled to reduce this error (Ma et al., 2020), but a more accurate estimation of water-use efficiency would include direct measurements of gas exchange and mesophyll conductance. Indeed, our observations are consistent with previous experiments in which the timing of leaf emergence, under low and high water availability, shifted both δ^13^C ratios and actual A/gs to a greater degree than the drought treatment (Franks, 2011). Finally, not all species were present at all sites (table S1), and therefore species identity may be confounded with environmental conditions. To address this, we looked at the effects of control and drought treatments which are paired at each site. All functional groups were represented at all sites, which allowed for a cross-site comparison of structure-function relationships. The best model was a global one (table S3), which included all functional groups across sites. Our analysis incorporates species, treatment, and site effects, whose relative predictive power can be parsed statistically (Table 2) using plot random effects and MPD to evaluate covariation between plots. Our data were plant-centric and focused on aboveground traits. However, belowground traits and microbial interactions could further improve models to predict form-function relationships at community and ecosystem levels (Silva & Lambers, 2021).

### 4.5 Conclusion

We found that the experimental drought stress did not affect leaf trait and stochiometric relationships across species. We used a mixed-effects model to infer variation in iWUE within and across 12 different species as a function of structural (SLA, life history strategy) and mean phylogenetic distance across species and sites. Our analyses included all relevant functional groups found in managed and natural Pacific Northwest grasslands and the results were consistent along a climate gradient from central Washington to southern Oregon. We found that species-specific responses were important across functional groups, a factor that should be taken into consideration for future management and restoration decisions. Going forward, our findings could inform trait-based approaches to predict water and nutrient use in ecosystems restoration and climate change mitigation and adaptation projects in the Pacific Northwest and other temperate grassland systems.

## Acknowledgements

The authors thank the Siskiyou Field Institute, The Nature Conservancy, and Capitol Land Trust for providing sites for this experiment, Laurel Pfeifer-Meister, Bitty Roy, Bart Johnson, Graham Bailes, Aaron Nelson, and Matthew Krna for their contributions to experimental design, and numerous others for assistance with the HOPS project. This experiment was funded by National Science Foundation Macrosystems Biology grant #1340847, Plant Biotic Interactions grant #1758947, and Convergence Accelerator Pilot grant #1939511.

## Author Contributions

All authors designed the research; HRD and TMM collected data; HRD analyzed data and wrote the paper; all authors edited the paper and helped interpret the data; LCRS funded the project.

## Supporting Information

Additional supporting information may be found online in the Supporting Information section at the end of the article.

Appendix 1: Supplemental tables and figures

## Open Research statement

The data used for all analyses in this study are available at the Dryad digital data repository via https://doi.org/10.5061/dryad.vmcvdncr7 with a CC0 1.0 license.

## Notes

https://doi.org/10.5061/dryad.vmcvdncr7

### Competing Interest Statement

The authors have declared no competing interest.

## References

Avise, J. C., & Wollenberg, K. (2010). Phylogenetics and the origin of species. Molecular Ecology and Evolution: The Organismal Side: Selected Writings from the Avise Laboratory, 94(July), 457–464. https://doi.org/10.1142/9789814317764_0007

Beer, C., Ciais, P., Reichstein, M., Baldocchi, D., Law, B. E., Papale, D., et al. (2009). Temporal and among-site variability of inherent water use efficiency at the ecosystem level. Global Biogeochemical Cycles, 23(2). https://doi.org/10.1029/2008GB003233

Blomberg, S. P., & Garland, T. (2002). Tempo and mode in evolution: Phylogenetic inertia, adaptation and comparative methods. Journal of Evolutionary Biology, 15(6), 899–910. https://doi.org/10.1046/j.1420-9101.2002.00472.x

Chase, M. W., Christenhusz, M. J. M., Fay, M. F., Byng, J. W., Judd, W. S., Soltis, D. E., et al. (2016). An update of the Angiosperm Phylogeny Group classification for the orders and families of flowering plants: APG IV. Botanical Journal of the Linnean Society, 181(1), 1–20. https://doi.org/10.1111/boj.12385

Clark, D. L., Wilson, M., Roberts, R., Dunwiddie, P. W., Stanley, A., & Kaye, T. N. (2012). Plant traits - a tool for restoration? Applied Vegetation Science, 15(4), 449–458. https://doi.org/10.1111/j.1654-109X.2012.01198.x

Dalton, M., & Fleishman, E. (2021). Fifth Oregon Climate Assessment. Corvallis, Oregon.

Enquist, B. J., Norberg, J., Bonser, S. P., Violle, C., Webb, C. T., Henderson, A., et al. (2015). Scaling from Traits to Ecosystems: Developing a General Trait Driver Theory via Integrating Trait-Based and Metabolic Scaling Theories. Advances in Ecological Research (1st ed., Vol. 52). Elsevier Ltd. https://doi.org/10.1016/bs.aecr.2015.02.001

Enquist, B. J., Bentley, L. P., Shenkin, A., Maitner, B., Savage, V., Michaletz, S., et al. (2017). Assessing trait-based scaling theory in tropical forests spanning a broad temperature gradient. Global Ecology and Biogeography, 26(12), 1357–1373. https://doi.org/10.1111/geb.12645

Franks, S. J. (2011). Plasticity and evolution in drought avoidance and escape in the annual plant Brassica rapa. New Phytologist, 190, 249–257. https://doi.org/10.1111/j.1469-8137.2010.03603.x

Gee, G., & Bauder, J. (1986). Particle-size analysis. In: Methods of Soil Analysis. In A. Klute (Ed.), Part 1. Physical and Mineralogical Methods (pp. 383–411). Madison, Wisconsin, USA: American Society of Agronomy.

Gouveia, A. C., & Freitas, H. (2009). Modulation of leaf attributes and water use efficiency in Quercus suber along a rainfall gradient. Trees - Structure and Function, 23(2), 267–275. https://doi.org/10.1007/s00468-008-0274-z

Hallett, L. M., Hsu, J. S., Cleland, E. E., Collins, S. L., Dickson, T. L., Farrer, E. C., et al. (2014). Biotic mechanisms of community stability shift along a precipitation gradient. Ecology, 95(6), 1693–1700. https://doi.org/10.1890/13-0895.1

Jackson, R. D., Paine, L. K., & Woodis, J. E. (2010). Persistence of native C4 grasses under high-intensity, short-duration summer bison grazing in the eastern tallgrass prairie. Restoration Ecology, 18(1), 65–73. https://doi.org/10.1111/j.1526-100X.2008.00439.x

Jin, Y., & Qian, H. (2019). V.PhyloMaker: an R package that can generate very large phylogenies for vascular plants. Ecography, 42(8), 1353–1359. https://doi.org/10.1111/ecog.04434

Jung, I. W., & Chang, H. (2012). Climate change impacts on spatial patterns in drought risk in the Willamette River Basin, Oregon, USA. Theoretical and Applied Climatology, 108(3–4), 355–371. https://doi.org/10.1007/s00704-011-0531-8

Katabuchi, M., & Masatoshi, K. (2015). LeafArea: an R package for rapid digital image analysis of leaf area. Ecological Research, 30(6), 1073–1077. https://doi.org/10.1007/s11284-015-1307-x

Keeling, R. F., Piper, S. C., Bollenbacher, A. F., & Walker, S. J. (2010). Monthly atmospheric 13C/12C isotopic ratios for 11 SIO stations. In Trends: A Compendium of Data on Global Change. Oak Ridge, Tenn., U.S.A.

Kröber, W., Plath, I., Heklau, H., & Bruelheide, H. (2015). Relating stomatal conductance to leaf functional traits. Journal of Visualized Experiments, 2015(104). https://doi.org/10.3791/52738

Li, D., Miller, J. E. D., & Harrison, S. (2019). Climate drives loss of phylogenetic diversity in a grassland community. Proceedings of the National Academy of Sciences of the United States of America, 116(40), 19989–19994. https://doi.org/10.1073/pnas.1912247116

Li, Z., Yang, L., Lu, W., Guo, W., Gong, X., Xu, J., & Yu, D. (2015). Spatial patterns of leaf carbon, nitrogen stoichiometry and stable carbon isotope composition of Ranunculus natans C.A. Mey. (Ranunculaceae) in the arid zone of northwest China. Ecological Engineering, 77, 9–17. https://doi.org/10.1016/j.ecoleng.2015.01.010

Ma, W. T., Tcherkez, G., Wang, X. M., Schäufele, R., Schnyder, H., Yang, Y., & Gong, X. Y. (2020). Accounting for mesophyll conductance substantially improves 13 C-based estimates of intrinsic water-use efficiency. New Phytologist, 0. https://doi.org/10.1111/nph.16958

Mackie, K. A., Zeiter, M., Bloor, J. M. G., & Stampfli, A. (2019). Plant functional groups mediate drought resistance and recovery in a multisite grassland experiment. Journal of Ecology, 107(2), 937–949. https://doi.org/10.1111/1365-2745.13102

Maxwell, T. M., Silva, L. C. R., & Horwath, W. R. (2018). Integrating effects of species composition and soil properties to predict shifts in montane forest carbon–water relations. Proceedings of the National Academy of Sciences of the United States of America, 115(18), E4219–E4226. https://doi.org/10.1073/pnas.1718864115

Mazerolle, M. J. (2006). Improving data analysis in herpetology: Using Akaike’s information criterion (AIC) to assess the strength of biological hypotheses. Amphibia Reptilia, 27(2), 169–180. https://doi.org/10.1163/156853806777239922

Mazerolle, M. J. (2020). AICcmodavg: Model selection and multimodel inference based on (Q)AIC(c).

Myers, S. S., Zanobetti, A., Kloog, I., Huybers, P., Leakey, A. D. B., Bloom, A. J., et al. (2014). Increasing CO2 threatens human nutrition. Nature, 510(7503), 139–142. https://doi.org/10.1038/nature13179

Ness, J. H., Rollinson, E. J., & Whitney, K. D. (2011). Phylogenetic distance can predict susceptibility to attack by natural enemies. Oikos, 120(9), 1327–1334. https://doi.org/10.1111/j.1600-0706.2011.19119.x

Ocheltree, T. W., Mueller, K. M., Chesus, K., LeCain, D. R., Kray, J. A., & Blumenthal, D. M. (2020). Identification of suites of traits that explains drought resistance and phenological patterns of plants in a semi-arid grassland community. Oecologia, 192(1), 55–66. https://doi.org/10.1007/s00442-019-04567-x

Ooms, J. (2019). magick: Advanced Graphics and Image-Processing in R.

Osborne, C. P., Salomaa, A., Kluyver, T. A., Visser, V., Kellogg, E. A., Morrone, O., et al. (2014). A global database of C4 photosynthesis in grasses. New Phytologist, 204, 441–446. https://doi.org/10.1111/nph.12942

Perez, T. M., & Heberling, M. (2020). Herbarium-based measurements reliably estimate three functional traits Plant invasions View project. https://doi.org/10.1002/ajb2.1535

Peterson, M. L., Bailes, G., Hendricks, L. B., Pfeifer-Meister, L., Reed, P. B., Bridgham, S. D., et al. (2021). Latitudinal gradients in population growth do not reflect demographic responses to climate. Ecological Applications, 31(2). https://doi.org/10.1002/EAP.2242

R Development Core Team. (2017). A Language and Environment for Statistical Computing. R Foundation for Statistical Computing. Vienna, Austria: R Foundation for Statistical Computing.

Reed, P. B., Pfeifer-Meister, L. E., Roy, B. A., Johnson, B. R., Bailes, G. T., Nelson, A. A., et al. (2019). Prairie plant phenology driven more by temperature than moisture in climate manipulations across a latitudinal gradient in the Pacific Northwest, USA. Ecology and Evolution, 9(6), 3637–3650. https://doi.org/10.1002/ece3.4995

Reed, P. B., Bridgham, S. D., Pfeifer-Meister, L. E., DeMarche, M. L., Johnson, B. R., Roy, B. A., et al. (2021). Climate warming threatens the persistence of a community of disturbance-adapted native annual plants. Ecology, 102(10), e03464. https://doi.org/10.1002/ecy.3464

Reed, P. B., Pfeifer-Meister, L. E., Roy, B. A., Johnson, B. R., Bailes, G. T., Nelson, A. A., & Bridgham, S. D. (2021). Introduced annuals mediate climate-driven community change in Mediterranean prairies of the Pacific Northwest, USA. Diversity and Distributions, 00, 1–12. https://doi.org/10.1111/ddi.13426

Rueden, C. T., Schindelin, J., Hiner, M. C., DeZonia, B. E., Walter, A. E., Arena, E. T., & Eliceiri, K. W. (2017). ImageJ2: ImageJ for the next generation of scientific image data. BMC Bioinformatics, 18(1), 529. https://doi.org/10.1186/s12859-017-1934-z

Saxton, K. E., & Rawls, W. J. (2006). Soil Water Characteristic Estimates by Texture and Organic Matter for Hydrologic Solutions. Soil Science Society of America Journal, 70(5), 1569–1578. https://doi.org/10.2136/SSSAJ2005.0117

Silva, L. C. R., & Lambers, H. (2021). Soil-plant-atmosphere interactions: structure, function, and predictive scaling for climate change mitigation. Plant and Soil, 1–23. https://doi.org/10.1007/s11104-020-04427-1

Silva, L. C. R., Pedroso, G., Doane, T. A., Mukome, F. N. D., & Horwath, W. R. (2015). Beyond the cellulose: Oxygen isotope composition of plant lipids as a proxy for terrestrial water balance. Geochemical Perspective Letters, 1, 33–42.

Soh, W. K., Yiotis, C., Murray, M., Parnell, A., Wright, I. J., Spicer, R. A., et al. (2019). Rising CO2 drives divergence in water use efficiency of evergreen and deciduous plants. Science Advances, 5(12), eaax7906. https://doi.org/10.1126/sciadv.aax7906

Temme, A. A., Liu, J. C., van Hal, J., Cornwell, W. K., Cornelissen, J. (Hans) H. C., & Aerts, R. (2017). Increases in CO2 from past low to future high levels result in “slower” strategies on the leaf economic spectrum. Perspectives in Plant Ecology, Evolution and Systematics, 29(September 2016), 41–50. https://doi.org/10.1016/j.ppees.2017.11.003

Tilman, D., & El Haddi, A. (1992). Drought and biodiversity in Grasslands. Oecologia, 89, 257–264.

USDA, & NRCS. (2019). The PLANTS Database (http://plants.usda.gov). Greensboro, NC: National Plant Data Team.

De Vienne, D. M., Aguileta, G., & Ollier, S. (2011). Euclidean nature of phylogenetic distance matrices. Systematic Biology, 60(6), 826–832. https://doi.org/10.1093/sysbio/syr066

Werner, C., Schnyder, H., Cuntz, M., Keitel, C., Zeeman, M. J., Dawson, T. E., et al. (2012). Progress and challenges in using stable isotopes to trace plant carbon and water relations across scales. Biogeosciences, 9(8), 3083–3111. https://doi.org/10.5194/bg-9-3083-2012

Van Wilgen, N. J., & Richardson, D. M. (2011). Is phylogenetic relatedness to native species important for the establishment of reptiles introduced to California and Florida? Diversity and Distributions, 17(1), 172–181. https://doi.org/10.1111/j.1472-4642.2010.00717.x

Wright, I. J., Reich, P. B., Westoby, M., Ackerly, D. D., Baruch, Z., Bongers, F., et al. (2004). The worldwide leaf economics spectrum. Nature, 428(6985), 821–827. https://doi.org/10.1038/nature02403

Yahdjian, L., & Sala, O. E. (2002). A rainout shelter design for intercepting different amounts of rainfall. Oecologia, 133(2), 95–101. https://doi.org/10.1007/s00442-002-1024-3

## References From the Supporting Information

Chaudhary, V. B., M. A. Rúa, A. Antoninka, J. D. Bever, J. Cannon, A. Craig, J. Duchicela, A. Frame, M. Gardes, C. Gehring, M. Ha, M. Hart, J. Hopkins, B. Ji, N. C. Johnson, W. Kaonongbua, J. Karst, R. T. Koide, L. J. Lamit, J. Meadow, B. G. Milligan, J. C. Moore, T. H. P. Iv, B. Piculell, B. Ramsby, S. Simard, S. Shrestha, J. Umbanhowar, W. Viechtbauer, L. Walters, G. W. T. Wilson, P. C. Zee, and J. D. Hoeksema. 2016. Data Descriptor: MycoDB, a global database of plant response to mycorrhizal fungi. Nature:3:160028.

Dickie, I. A., L. B. Martínez-García, N. Koele, G. A. Grelet, J. M. Tylianakis, D. A. Peltzer, and S. J. Richardson. 2013. Mycorrhizas and mycorrhizal fungal communities throughout ecosystem development. Plant and Soil 367:11–39.

Schloerke, B., D. Cook, J. Larmarange, F. Briatte, M. Marbach, E. Thoen, A. Elberg, and J. Crowley. 2020. GGally: Extension to “ggplot2.”

Wang, B., and Y. L. Qiu. 2006. Phylogenetic distribution and evolution of mycorrhizas in land plants. Mycorrhiza 16:299–363.

