## Supplementary material for "Leaf traits predict water-use efficiency in U.S. Pacific Northwest grasslands under rain exclusion treatment": SI Appendix 1

### Contents of this file

Fig. S1. Daily variation in effect of rain exclusion treatment on soil matric potential (SMP) at each site.
Fig. S2 Daily variation in effects of rain exclusion treatment on soil temperature at each site.
Fig. S3 Principal component analysis of specific leaf area (SLA) and three functional traits.
Fig. S4 Correlation between the four traits measured including specific leaf area (SLA), intrinsic water-use efficiency (iWUE), carbon to nitrogen ratio (C:N), and nitrogen (N).

Table S1. Complete list of species included in this study, replicates of each species, and plant strategies.
Table S2 Distribution of functional groups and life history strategies across sites.
Table S3 AICc values used in intrinsic water-use efficiency (iWUE) model selection.
Table S4 Statistical difference of rain exclusion treatment on soil matric potential.
Table S5 Statistical difference of rain exclusion treatment on soil temperature.
Table S6 Summary statistics of effects of rain exclusion treatment on soil matric potential (SMP).
Table S7. Summary statistics of effects of rain exclusion treatment on soil temperature.
Table S8. Statistical significance of each factor influencing intrinsic water-use efficiency (iWUE) model shown in Fig. 4.

Appendix 1 references

### Introduction

Included in this SI are four figures and eight tables that supplement the text. All methods of calculation are included in the text. The data used in these analyses are publicly available in a data repository. Please check the webpage for this article for a link to the dataset.

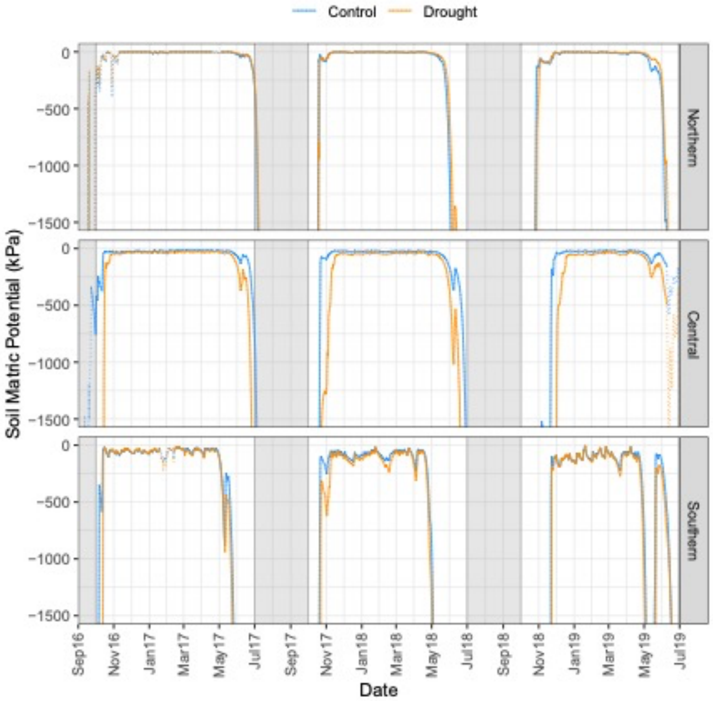

**Fig. S1. Daily variation in effect of rain exclusion treatment on soil matric potential (SMP) at each site.**

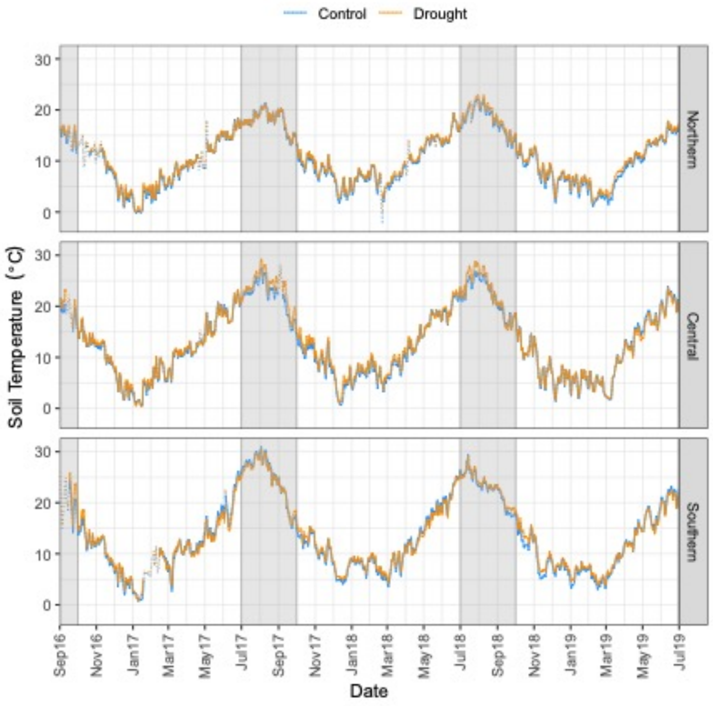

**Fig. S2 Daily variation in effects of rain exclusion treatment on soil temperature at each site**.

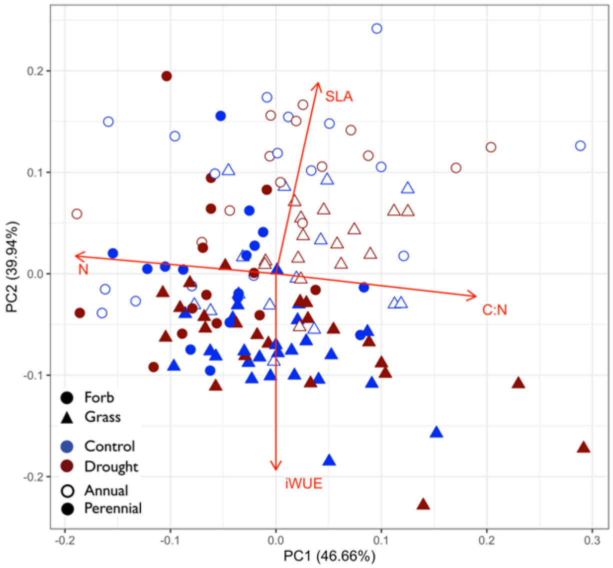

**Fig. S3 Principal component analysis of specific leaf area (SLA) and three functional traits.** SLA has a nearly orthogonal relationship with N content (N), intrinsic water-use efficiency (iWUE), and carbon to nitrogen ratio (C:N). Life history strategy (annual, perennial) is divided along PC2 while functional group (grass, forb) is divided along PC1. Model explained 86.6% of variation.

B

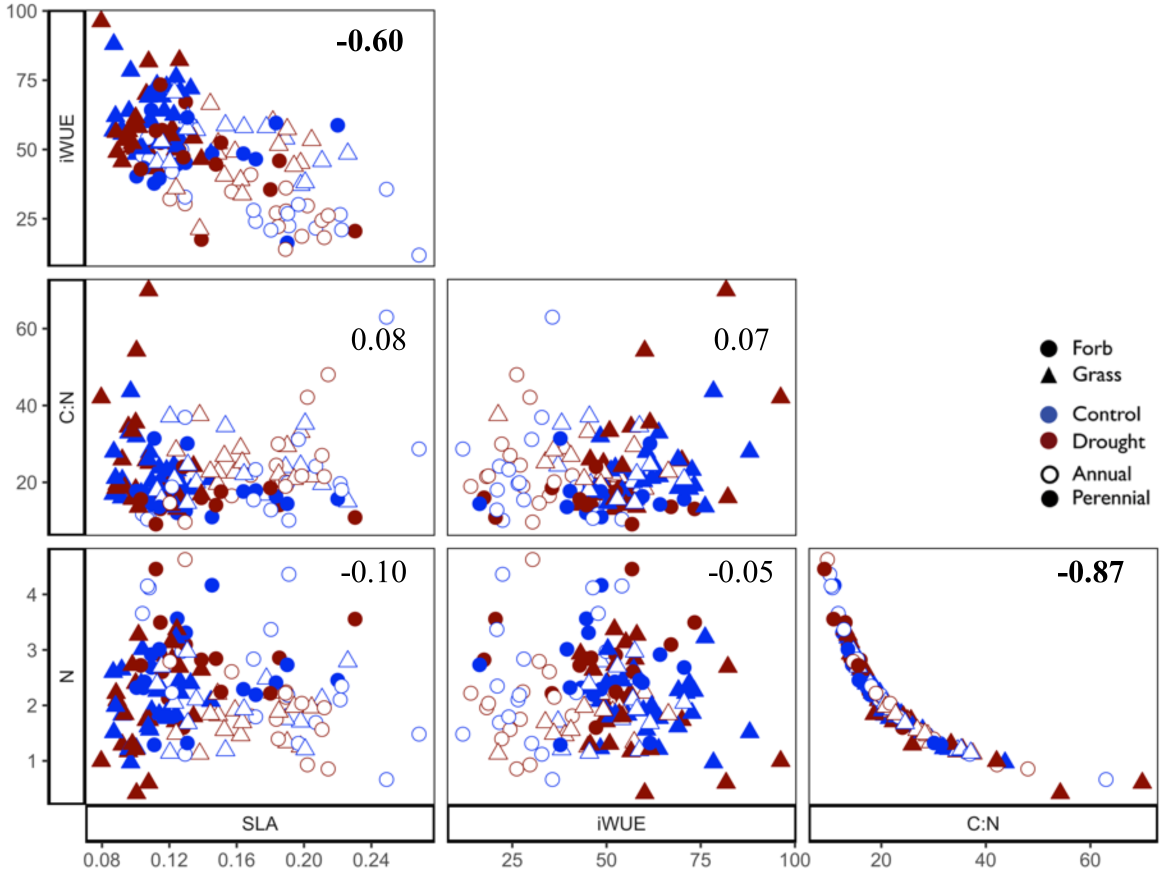

**Fig. S4 Correlation between the four traits measured including specific leaf area (SLA), intrinsic water-use efficiency (iWUE), carbon to nitrogen ratio (C:N), and nitrogen (N).** Each point represents a single species in each plot. Values shown are $R^2$, significance indicated in bold. Plot made with GGally package in R (Schloerke et al. 2020).

**Table S1. Complete list of species included in this study, replicates of each species, and plant strategies.**

| Species name | Family | Life strategy | Functional group | Total | Number sampled (SLA replicates) | | | Nutrient-use strategy |
|---|---|---|---|---|---|---|---|---|
| | | | | | *Northern* | *Central* | *Southern* | |
| *Agrostis capillaris* L. | Poaceae | Perennial | Grass | 9 (16) | 0 | 9 (16) | 0 | AM[1] |
| *Alopecurus pratensis* L. | Poaceae | Perennial | Grass | 19 (50) | 19 (50) | 0 | 0 | AM[2] |
| *Bromus diandrus* Roth | Poaceae | Annual | Grass | 9 (26) | 0 | 9 (26) | 0 | AM[1]* |
| *Bromus hordeaceus* L. | Poaceae | Annual | Grass | 21 (50) | 5 (15) | 4 (12) | 12 (23) | AM[1]* |
| *Daucus carota* L. | Apiaceae | Annual | Forb | 2 (6) | 0 | 0 | 2 (6) | AM[1] |
| *Elymus repens* (L.) Gould | Poaceae | Perennial | Grass | 10 (31) | 0 | 0 | 10 (31) | AM[1]* |
| *Geranium dissectum* L. | Malvaceae | Annual | Forb | 13 (36) | 10 (28) | 3 (8) | 0 | AM[2] |
| *Lotus corniculatus* L. | Fabaceae | Perennial | Forb | 3 (9) | 3 (9) | 0 | 0 | AM, N-fixer[1] |
| *Schedonorus arundinaceus* (Schreb.) Dumort. | Poaceae | Perennial | Grass | 10 (23) | 0 | 0 | 10 (23) | AM[1] |
| *Sidalcea malviflora* (DC.) A. Gray ex Benth. | Malvaceae | Perennial | Forb | 27 (81) | 10 (30) | 8 (23) | 9 (28) | AM[3] |
| *Trifolium subterraneum* L. | Fabaceae | Annual | Forb | 11 (34) | 0 | 9 (31) | 2 (3) | AM, N-fixer[1] |
| *Vicia sativa* L. | Fabaceae | Annual | Forb | 5 (15) | 5 (15) | 0 | 0 | AM, N-fixer[1]* |
| ***Totals*** | | | | **139 (368) 12 species** | **52 (138) 6 species** | **42 (116) 6 species** | **45 (114) 6 species** | |

Notes: Nutrient-use strategy data derived from [1]Chaudhary et al. (2016), [2]Wang and Qui (2006), and [3] Dickie et al. (2013). * indicates "to genus," that strategy was extrapolated from a sister species given a lack of data on this species. Nutrient-use strategies include arbuscular mycorrhizal (AM) and nitrogen fixation (N-fixer). Totals are given followed by the number of specific leaf area (SLA) replicates in parentheses.

### Table S2 Distribution of functional groups and life history strategies across sites.

| | Annual | | Perennial | |
|---|---|---|---|---|
| **Site** | **Grass** | **Forb** | **Grass** | **Forb** |
| Northern | 1 | 2 | 1 | 2 |
| Central | 2 | 2 | 1 | 1 |
| Southern | 1 | 2 | 2 | 1 |

Note: Counts represent individual species from each group sampled at each site.

### Table S3 AICc values used in intrinsic water-use efficiency (iWUE) model selection.

| | **Model** | **K** | **AICc** | **ΔAICc** | **Model Likelihood** | **AICc weights** | **Log likelihood** | **Cumulative AIC weights** |
|---|---|---|---|---|---|---|---|---|
| **Model 4** | SLA+MPD+C:N+N+ life history strategy | 11 | 1086.406 | 0.000 | 1.000 | 0.948 | -531.164 | 0.948 |
| **Model 2** | SLA+MPD | 8 | 1092.723 | 6.317 | 0.042 | 0.040 | -537.808 | 0.988 |
| **Model 3** | SLA+C:N+N+MPD | 10 | 1096.007 | 9.601 | 0.008 | 0.008 | -537.144 | 0.996 |
| **Model 5** | MPD+life history strategy | 8 | 1097.425 | 11.019 | 0.004 | 0.004 | -540.159 | 1.000 |
| **Model 7** | SLA+C:N+N+life history strategy | 10 | 1109.732 | 23.326 | 0.000 | 0.000 | -544.007 | 1.000 |
| **Model 1** | SLA | 7 | 1111.642 | 25.236 | 0.000 | 0.000 | -548.394 | 1.000 |
| **Model 6** | SLA+C:N+N | 9 | 1112.294 | 25.888 | 0.000 | 0.000 | -546.449 | 1.000 |

Note: All models include plot as random effect and rain exclusion treatment and site as a fixed effects. Models were selected using biologically likely hypotheses. Variables include specific leaf area (SLA), mean phylogenetic distance (MPD), C:N, N content, and life history strategy (annual, perennial).

**Table S4 Statistical difference of rain exclusion treatment on soil matric potential**.

| Year | Predictor | *F* | *P* |
|---|---|---|---|
| 2017 | Site | 179.79 | **< 0.001** |
| | Rain excl. trt | 3.98 | *0.057* |
| | Site x Rain excl. trt | 2.70 | *0.087* |
| 2018 | Site | 146.40 | **< 0.001** |
| | Rain excl. trt | 5.79 | **0.024** |
| | Site x Rain excl. trt | 2.16 | 0.137 |
| 2019 | Site | 91.59 | **< 0.001** |
| | Rain excl. trt | 4.12 | *0.054* |
| | Site x Rain excl. trt | 4.42 | **0.023** |

Notes: Significance tested with an ANOVA. Growing season is defined as 1 Oct to 30 Jun and each season is labeled by the spring months of the growing year (e.g. 2018 is 1 Oct 2017 to 30 Jun 2018).

**Table S5 Statistical difference of rain exclusion treatment on soil temperature**.

| Year | Predictor | *F* | *P* |
|---|---|---|---|
| 2017 | Site | 216.97 | **< 0.001** |
| | Rain excl. trt | 4.58 | **0.043** |
| | Site x Rain excl. trt | 0.91 | 0.417 |
| 2018 | Site | 230.89 | **< 0.001** |
| | Rain excl. trt | 15.94 | **0.001** |
| | Site x Rain excl. trt | 1.52 | 0.239 |
| 2019 | Site | 168.30 | **< 0.001** |
| | Rain excl. trt | 5.45 | **0.028** |
| | Site x Rain excl. trt | 3.32 | *0.053* |

Notes: Significance tested with an ANOVA. Growing season is defined as 1 Oct to 30 Jun and each season is labeled by the spring months of the growing year (e.g. 2018 is 1 Oct 2017 to 30 Jun 2018).

**Table S6 Summary statistics of effects of rain exclusion treatment on soil matric potential (SMP).**

| Year | Site | Treatment | N | T-ratio | *P* | mean SMP | SD |
|---|---|---|---|---|---|---|---|
| 2017 | Southern | Control | 5 | 0.702 | 0.490 | -90.803 | 47.501 |
| | | Drought | 5 | | | -114.127 | 68.523 |
| | Central | Control | 5 | 2.975 | **0.007** | -27.233 | 1.863 |
| | | Drought | 5 | | | -54.831 | 4.762 |
| | Northern | Control | 5 | -0.220 | 0.828 | -4.057 | 1.953 |
| | | Drought | 5 | | | -3.554 | 0.409 |
| 2018 | Southern | Control | 5 | 1.163 | 0.256 | -237.932 | 111.111 |
| | | Drought | 5 | | | -356.978 | 205.041 |
| | Central | Control | 5 | 2.958 | **0.007** | -44.947 | 6.258 |
| | | Drought | 5 | | | -97.448 | 14.419 |
| | Northern | Control | 5 | 0.045 | 0.964 | -11.435 | 5.276 |
| | | Drought | 5 | | | -10.864 | 2.540 |
| 2019 | Southern | Control | 5 | 0.662 | 0.514 | -312.293 | 189.753 |
| | | Drought | 5 | | | -393.405 | 245.943 |
| | Central | Control | 5 | 3.481 | **0.002** | -66.543 | 13.957 |
| | | Drought | 5 | | | -174.131 | 31.206 |
| | Northern | Control | 5 | -0.629 | 0.535 | -24.686 | 8.689 |
| | | Drought | 5 | | | -20.215 | 6.247 |

Notes: Growing season is defined as 1 Oct to 30 Jun and each season is labeled by the spring months of the growing year (e.g. 2018 is 1 Oct 2017 to 30 Jun 2018). T-ratios and P-values are for a Tukey post hoc test on the ANOVA results presented in Table S4.

**Table S7. Summary statistics of effects of rain exclusion treatment on soil temperature.**

| Year | Site | Treatment | N | T-ratio | *P* | Mean soil temp | Sd |
|---|---|---|---|---|---|---|---|
| 2017 | Southern | Control | 5 | | | 10.974 | 0.271 |
| | | Drought | 5 | -0.37 | 0.714 | 11.033 | 0.248 |
| | Central | Control | 5 | | | 10.787 | 0.251 |
| | | Drought | 5 | -1.08 | 0.293 | 10.958 | 0.373 |
| | Northern | Control | 5 | | | 8.736 | 0.113 |
| | | Drought | 5 | -2.26 | **0.033** | 9.094 | 0.168 |
| 2018 | Southern | Control | 5 | | | 11.615 | 0.263 |
| | | Drought | 5 | -0.97 | 0.344 | 11.779 | 0.219 |
| | Central | Control | 5 | | | 10.683 | 0.099 |
| | | Drought | 5 | -3.39 | **0.002** | 11.261 | 0.505 |
| | Northern | Control | 5 | | | 8.968 | 0.104 |
| | | Drought | 5 | -2.55 | **0.017** | 9.402 | 0.206 |
| 2019 | Southern | Control | 5 | | | 11.018 | 0.233 |
| | | Drought | 5 | -1.56 | 0.131 | 11.306 | 0.260 |
| | Central | Control | 5 | | | 10.583 | 0.239 |
| | | Drought | 5 | 0.57 | 0.573 | 10.478 | 0.448 |
| | Northern | Control | 5 | | | 8.570 | 0.279 |
| | | Drought | 5 | -3.05 | **0.005** | 9.132 | 0.226 |

Notes: Growing season is defined as 1 Oct to 30 Jun and each season is labeled by the spring months of the growing year (e.g. 2018 is 1 Oct 2017 to 30 Jun 2018). T-ratios and P-values are for a Tukey post hoc test on the ANOVA results presented in Table S5.

**Table S8. Statistical significance of each factor influencing intrinsic water-use efficiency (iWUE) model shown in Fig. 4.**

| **Predictor** | $\chi^2$ | *P* |
|---|---|---|
| *Fixed effects* | | |
| Specific leaf area | 13.869 | **<0.001** |
| Mean phylogenetic distance | 28.212 | **<0.001** |
| C:N | 2.726 | *0.099* |
| N | 1.435 | 0.231 |
| Life history strategy (annual/perennial) | 12.491 | **<0.001** |
| Treatment | 4.938 | **0.026** |
| Site | 2.745 | 0.254 |
| *Random effect* | | |
| Plot | | 1 |

Note: Significance tested with a Type II ANOVA.
